# Mesoscopic SCAPE Microscope with a Rescanned, Super-oblique Illumination Plane

**DOI:** 10.1101/2025.04.25.650616

**Authors:** Zixian Cao, Jiapeng Zhu, Cheng Zhang, Qianqian Wang, Yankan Huang, Wei Liu, Bingxin Shen, Yuming Chai, Zhaomin Zhong, Li He, Quan Wen, Han Wang, Wenxuan Liang

## Abstract

Investigating structural and functional dynamics across large spatial scales in living organisms calls for volumetric microscopy that combines a wide field of view (FOV), high spatial resolution, and sufficient temporal resolution. Swept confocally-aligned planar excitation (SCAPE) microscopy offers a compelling balance of optical sectioning, cellular resolution, and high-speed volumetric imaging. However, extending SCAPE to mesoscopic scales has been hindered by the limited tilting angle of the oblique illumination light sheet achievable with commercial low-magnification objectives. Here, we present a mesoscopic SCAPE microscope that overcomes this constraint by implementing a super-oblique light sheet—tilted up to ~60° relative to the optical axis of a 0.5-NA primary objective—enabled by an air-to-water bridging reflector. In addition, we introduce a single-galvanometer, self-conjugating rescanning strategy that allows arbitrary customization of the optical-to-mechanical angle ratio. This innovation accommodates the reversed and modulated scan displacement of the reflected light sheet and preserves descanned epi-fluorescence detection. Together, these advancements enable our mesoscopic SCAPE microscope to achieve a lateral FOV of 5.0 mm × 2.9 mm, cellular-level spatial resolution (~9.4 μm × ~5.9 μm lateral, and ~6.0 μm axial), and a volumetric imaging rate of 9 volumes per second.

## 1. Introduction

Organisms and their environments are intrinsically three-dimensional (3D), necessitating high-speed depth-resolved volumetric microscopy to study their structural and functional dynamics over time. Even simple animal behaviors often involve coordinated activity across multiple brain regions, while more complex functions—such as learning, memory, and consciousness— require the integration of neurobiological processes spanning vast spatial and temporal scales. To decipher inter-neuron communication, it is essential to record the activity of hundreds of thousands of neurons simultaneously across large fields of view (FOV) and multiple depths, with high spatial and temporal resolution.

The development of calcium indicators (e.g., GCaMP) and transgenic animal models in recent decades has spurred the development of a plethora of *in vivo* fluorescence microscopy methods for large-FOV neuronal imaging [1]. While wide-field optical imaging can capture cortex-wide *in vivo* neural activity in mouse brains, it integrates signals from all depths, lacking both depth-resolving capability and cellular-level specificity [2, 3]. Conventional point-scanning modalities such as confocal and two-photon fluorescence microscopy achieve sub-cellular resolution and can cover multi-millimeter FOVs [4–8], but their volumetric imaging rate remains limited (typically on the order of ~1 volume/s) due to scanner inertia, phototoxicity and sample irritation concerns, and laser repetition rate constraints [9]. Light-sheet microscopy (LSM), or selective plane illumination microscopy (SPIM), overcomes some of these limitations through parallelized planar illumination and detection, affording faster volumetric imaging with reduced phototoxicity [10–12]; however, its orthogonal dual-objective geometry imposes significant sample space restrictions, largely confining its applications to cultured cells and developing embryos.

Swept confocally-aligned planar excitation (SCAPE) microscope [13, 14]—also referred to as epi-illumination selective plane illumination microscope (eSPIM) [15], scanned oblique plane microscope (OPM) [16–18] or scanned oblique plane illumination (SOPi) microscope [19]—achieves a well-balanced combination of optical sectioning capability, cellular resolution and volumetric imaging rate. Leveraging the ample aperture angle of a high-numerical aperture (NA) objective, SCAPE generates an oblique illumination light sheet while collecting epi-fluorescence through the same primary objective (O1); in this way, it affords an open sample space compatible with diverse specimens and animal models, including *C. elegans, Drosophila* larvae, fruit flies, larval zebrafish, and live mouse and human tissues [14, 20–23]. Another key innovation of SCAPE is its use of a galvanometer mirror conjugated to O1’s back focal plane, which simultaneously sweeps the oblique illumination plane and descans the epi-fluorescence. This creates a stationary intermediate image in the focal region of the secondary objective (O2), which is then relayed by a tertiary objective (O3) onto an sCMOS camera, thereby enabling translation-free, fast volumetric imaging. To date, reported SCAPE prototypes have generally achieved volumetric imaging rates on the order of 10 volume/sec, with the upper limit determined by sCMOS frame rate (on the order of 1000 fps in rolling-shutter mode).

To extend SCAPE microscopy to mesoscopic FOVs, one could, in principle, utilize custom-designed high-NA, large-field objectives [4–8]. However, commercially available low-magnification objectives offer more practical advantages in terms of cost-effectiveness, availability, and ease of integration. A fundamental limitation arises from the inherent trade-off between field size and aperture angle in objective design: commonly used commercial objectives that support multi-millimeter FOVs typically have NAs ≤ 0.5, corresponding to aperture angles up to 30° in air. This restricts both the oblique illumination angle and the intermediate image angle to below 30°. Consequently, to properly focus on the intermediate image, the optical axis of O3 must be tilted by more than 60° relative to O2. This leads to poor overlap between the acceptance cone of O3 and the emission cone of O2, which in turn compromises the effective aperture angle, spatial resolution, fluorescence collection efficiency, and ultimately, the volumetric imaging rate.

To address aforementioned challenges associated with the angle of the illumination sheet angle, past solutions can be broadly categorized in two technical approaches. One approach focuses on modifying the emission cone of the intermediate fluorescence image formed by O2, or adjusting the collection cone of O3, to increase their overlap. Representative strategies include projecting the intermediate image onto a blazed grating [24] or the high-acceptance NA endface of a fiber-optic faceplate [25], or tuning the system magnification to produce a minified intermediate image with a steeper oblique angle [26]. While these work-arounds significantly enhance collection efficiency, the spatial resolution—particularly the axial resolution— remains fundamentally constrained by the limited angle between the illumination light sheet and the effective detection point-spread function (PSF), both of which are bounded by the low aperture angle of the large-field, low-NA primary objective (O1).

In light of these limitations, a potentially more effective strategy is to generate an illumination light sheet with an oblique angle—relative to the axis of O1—that exceeds O1’s aperture angle, so as to enhance both collection efficiency and spatial resolution simultaneously.

One approach involves deploying a transmission grating within the working space of O1 to diffract and bend the light sheet [27]. Although relatively straightforward to implement, the grating generates higher-order diffracted light sheets that sweep along with the primary oblique light sheet and must be blocked dynamically, complicating the illumination path and risking occlusion of epi-fluorescence. Moreover, the grating diffracts the epi-fluorescence light itself, introducing aberrations and reducing collection efficiency. Another solution employs an auxiliary illumination objective positioned near—but at a non-orthogonal angle to—O1 to deliver an arbitrarily oblique light sheet into O1’s focal region [28]. However, this design necessitates an additional galvanometer mirror dedicated to scanning the illumination light sheet in synchrony with the descanning galvanometer in the detection path to maintain a stationary intermediate epi-fluorescence image. One recent approach involves guiding an initial illumination plane—positioned off-axis and approximately parallel to the optical axis of O1— to reflect off the oblique inner sidewall of a thick, trapezoid-shaped glass slide. Such configuration generates a so-called mirrored light sheet that enters the sample at a significantly steeper oblique angle [29]. The main drawback of this method is that this mirrored light sheet sweeps in the opposite direction—and at a different lateral translation speed—compared to the initial light sheet, disrupting the natural descanning mechanism of SCAPE. To restore descanned epi-fluorescence detection, the authors introduced a dual-galvo scanner unit into the illumination path, in addition to the primary galvo scanner (shared by both illumination and detection path), to compensate for the modulated lateral displacement of the mirrored light sheet, thereby increasing the system complexity.

Here, we present a single-galvanometer, single-primary objective, mesoscopic SCAPE microscope that leverages a water immersion-compatible reflector unit (or an air-to-water bridging reflector unit) to generate a super-oblique illumination plane tilted by approximately 60° in water (relative to the optical axis of O1); this significantly exceeds the aperture angle corresponding to O1’s 0.5 NA. Additionally, we introduce a single-galvanometer, self-conjugating rescanning strategy that allows flexible customization of the optical-to-mechanical angle ratio (OMAR), thereby accommodating the reversed scan of the reflected light-sheet and preserving epi-fluorescence descanning. In the following sections, we outline the design principles of rescanned, super-oblique illumination plane, characterize the resultant meso-SCAPE system, and demonstrate its capability for high-speed, mesoscale volumetric imaging in live animal models.

## 2. Design principles

The high volumetric imaging rate of SCAPE microscopy relies on two core principles: (1) oblique plane illumination that encodes depth into lateral position on the intermediate image, and (2) descanned epi-fluorescence detection which reconstructs a stationary intermediate image. To achieve a multi-millimeter FOV, we employ a commercially available low-magnification objective (MVPLAPO 2XC, Olympus) with an NA of 0.5 and a usable FOV of approximately 7.0 mm in diameter [30]. Leveraging this objective’s 20-mm-long working distance (WD), we designed and custom-built a trapezoidal prism from a refractive index (RI)-matched polymer (RI = 1.329 at 589 nm; BIO-133, MY Polymers) to bridge the air and water immersion environments of live animal models (Fig. 1). This configuration enables us to tilt the illumination plane beyond the objective’s aperture angle while minimizing aberrations to the light sheet caused by refraction at the air-water interface under high incidence angles. The initial illumination plane exits O1 and impinges the air-facing surface of the polymer prism perpendicularly, thereby minimizing refraction. It is then reflected by a sidewall inclined at *α* ≈ 30° relative to the optical axis of O1, passes through the polymer-water interface without refraction, and finally enters the biological tissue at an oblique angle of *β* ≈ 60° with minimal distortion. While this reflected light sheet design is conceptually related to the glass microprism approach in ref. [29], our polymer-based implementation allows a upright imaging geometry and facilitates straightforward access to live animal models.

**Fig. 1.**
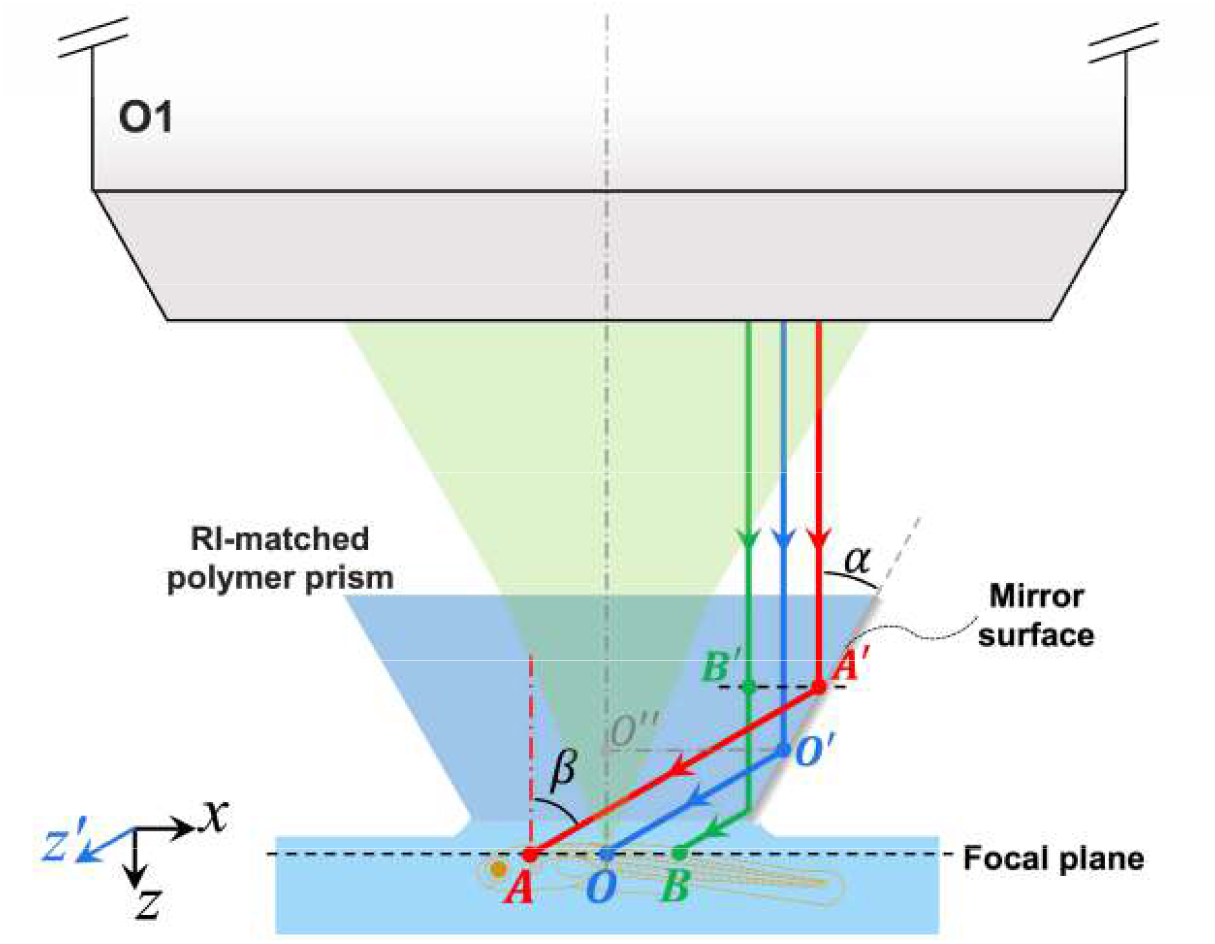
Schematic of the super-oblique light sheet generation using an air-to-water bridging reflector. A trapezoidal prism was fabricated from an ultraviolet-curable optical polymer with a refractive index matched to water; one angled sidewall was directly bonded to a silver-coated miniature mirror during fabrication to ensure uniform reflection of the incoming light sheet. The red, green, and blue lines indicate the chief rays of the light sheet in the *xz*-plane at different scan positions, with the light sheet’s width extending perpendicular to the plane of the page.

One prominent challenge associated with adopting this polymer trapezoidal prism lies in the reversed and modulated lateral motion of the reflected illumination plane compared to the pre-reflection light sheet. Specifically, from Fig. 1, the displacement of the reflected light sheet is given by 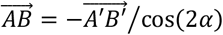, indicating both a reversal in direction and a change in scan velocity. Such changes disrupts the standard descanning mechanism of SCAPE and scanning OPM. To illustrate that, we draw in Fig. 2A a conceptual schematic of the illumination and detection beam paths, spanning from the galvanometer (galvo) mirror to the primary objective O1, with the bridging reflector positioned within the working space of O1. With the mechanical deflection angle of the galvo mirror (defined as positive in the counter-clockwise direction) denoted by *θ*_galvo_, the optical scan angle of illumination beam exiting the galvo mirror is simply *θ*_ill_ = 2*θ*_galvo_. Then after passing through O1, the illumination beam reflects off the angled sidewall of the polymer prism, and the resulting epi-fluorescence beam returns directly through O1 and eventually reflects off the deflected galvo mirror. The deflection angle of the returning epi-fluorescence beam equals (derived in detail in Supplementary Note S1)

**Fig. 2.**
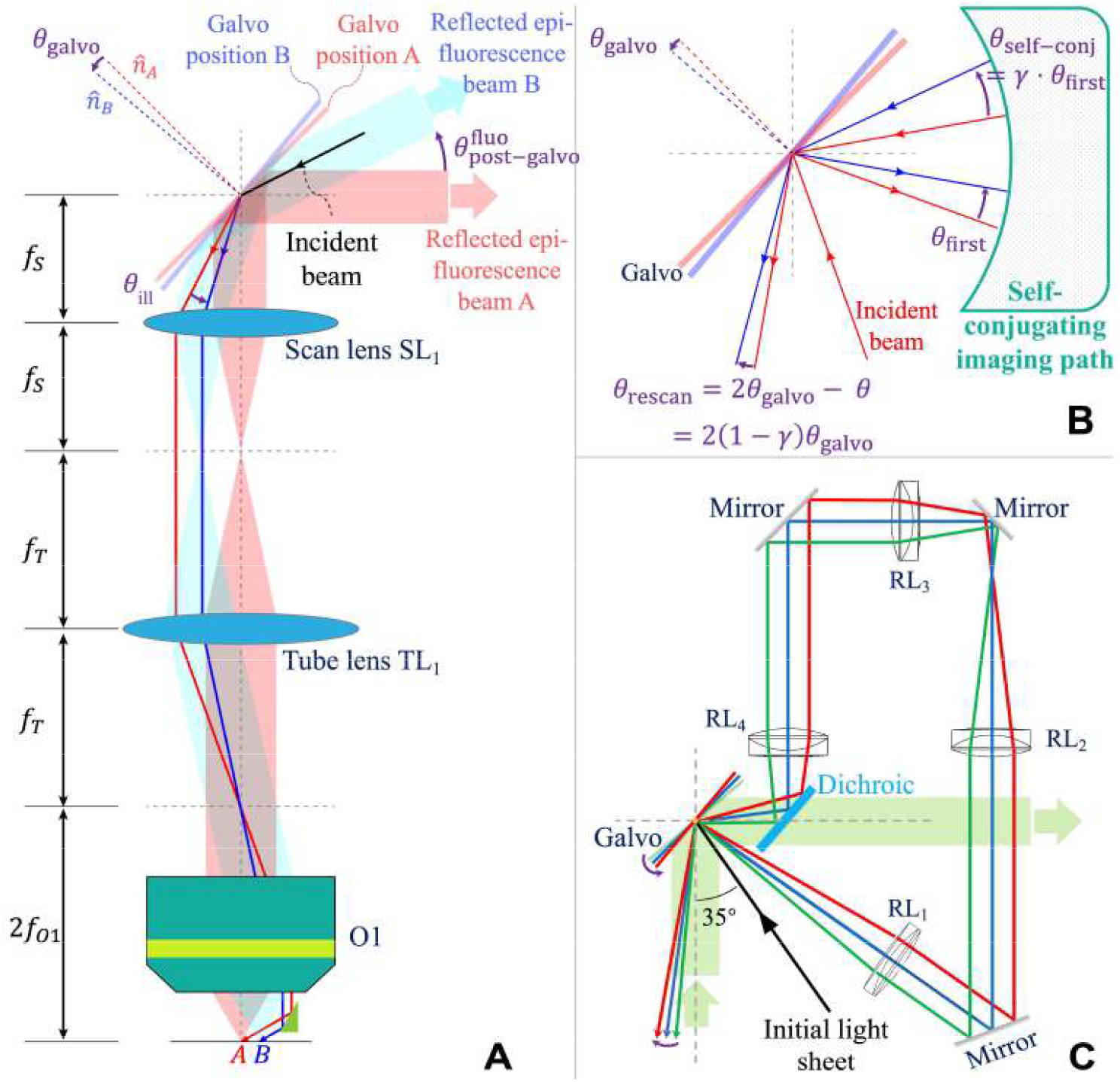
Self-conjugating rescanning strategy. A) When the reflector is introduced under O1, epi-fluorescence cannot be descanned. B) Self-conjugated rescan module can adjust the scanning of illumination light path. C) After the incident light is reflected by the galvanometer, it passes through an 8f system (GL_1_-GL_2_-GL_3_-GL_4_) and then returns to the galvanometer again. The excitation light path of the galvanometer at different positions is distinguished by different colors.

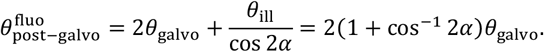

Clearly for our target angle of *α* ≈ 30°, the coefficient (1 + cos^-1^ 2*α*) ≠ 0, implying that the returning fluorescence beam is not de-scanned. Instead, it deflects along with the rotating galvo mirror.

To correct the altered lateral sweep velocity without introducing additional galvanometers, we developed a self-conjugating rescanning strategy that allows flexible customization of the optical-to-mechanical scan angle ratio (OMAR), defined as *b* ≐ *θ*_ill_/*θ*_galvo_. In this approach, the optical scan angle of the exiting illumination beam is no longer constrained to be twice the mechanical deflection angle of the galvanometer. The core concept behind achieving a customizable OMAR is to introduce an imaging path which maps the pivot point of the galvanometer back onto itself—hence the term “self-conjugating”. After its initial reflection by the galvanometer, the illumination beam is relayed through the imaging optical path, re-impinges onto the same galvanometer, and gets re-scanned by the rotated galvanometer mirror. This second interaction results in a rescanned illumination beam, whose optical scan angle can be described as (detailed in Supplementary Note S1)

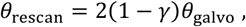

where γ denotes the angular magnification of the self-conjugating imaging path, resulting in an OMAR of *b* = 2(1 − γ). By designing the self-conjugating imaging path such that γ = 1 + cos 2*α*, we can tune the OMAR to be *b* = −2 cos 2*α*, and then

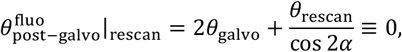

i.e. the returning fluorescence beam is descanned by the galvo.

Note the above analysis assumes the returning epi-fluorescence beam does not traverse the self-conjugating optical path, but instead impinges the galvanometer mirror and reflects only once. This can be readily realized by incorporating a dichroic mirror within the self-conjugating imaging path to transmit the returning epi-fluorescence beam. As illustrated in Fig. 2C, the imaging path implemented in our system consists of two concatenated 4f optical relays, resulting in an overall angular magnification of γ = 1.5 and matching the sidewall inclination angle α = 30° of the polymer prism. The collected epi-fluorescence beam, after being reflected and descanned by the galvanometer mirror, passes through the dichroic mirror and forms a stationary intermediate image in the focal region of O2.

Following the principles outlined above, we constructed a prototype meso-SCAPE microscope that features a super-oblique light-sheet inclined by 60 degrees in water. As illustrated in Fig. 3, the complete optical setup comprises five main modules (see Methods for details on design considerations and components used in each module):

**Fig. 3.**
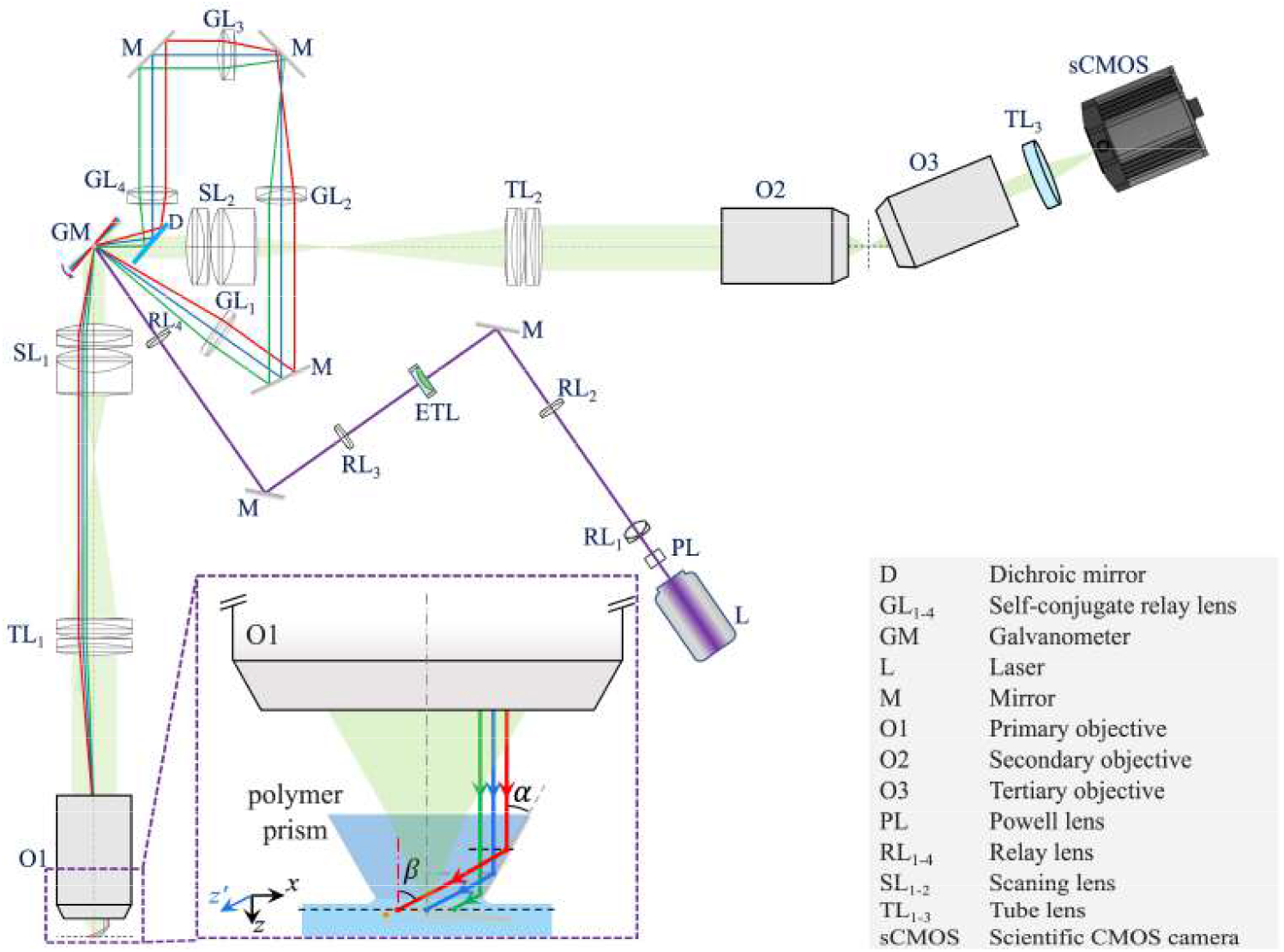
Schematic of the meso-SCAPE microscope. Essentially, the illumination light emitted from the laser (L) first passes a Powell lens (PL) to form an initial light sheet of improved illumination uniformity. It then travels through relay lenses RL_1_-RL_4_, an electrical tunable lens (ETL), and enters the scanning galvanometer (GM). The red-, green-, and blue-colored rays correspond to three different deflection angles of the galvanometer. The illumination light, after its initial reflection by the galvanometer, passes a self-conjugating optical path comprised of lenses GL_1_-GL_4_, re-enters the galvanometer, and then undergoes a second scan. The exiting illumination beam then passes through a scan lens SL_1_ and a tube lens TL_1_, before entering the primary objective lens O1 and the RI-matched polymer reflector to achieve an oblique angle of 60°. Epi-fluorescence originating from the sample space (light green shading) travels through the primary detection optical path (consisting of O1, SL_1_, TL_1_, GM, SL_2_, TL_2_, and O2 in sequence), gets descanned, and forms a stationary intermediate image in the focal region of O2, which is then collected by a tertiary objective lens O3, and relayed through a tube lens TL3 onto an sCMOS camera for recording. See Supplementary Note S3 for a photograph of the prototype microscope.

1. The initial light sheet generation module, which employs a Powell lens to generate the initial light sheet and an electrically tunable lens (ETL) to dynamically adjust the sheet waist.
2. The self-conjugating rescanning module, which scans the illumination sheet with a customized OMAR to ensure descanned epi-fluorescence detection.
3. The primary objective module, which includes O1, the associated scan lens (SL_1_) and tube lens (TL_1_), and the air-to-water bridging reflector, to generate the super-oblique illumination plane.
4. The secondary objective module, consisting of O2 (the same as O1), along with its scan lens (SL_2_) and tube lens (TL_2_), which forms a stationary intermediate image near the focus of O2.
5. The tertiary objective module, which includes O3 and a matching tube lens (TL_3_) and magnifies the intermediate image onto the sCMOS camera for data acquisition.

Note that while in principle total internal reflection could be leveraged to reflect the light sheet off the inner sidewall of the polymer prism, in practice we built the polymer prism around a silver-coated miniature mirror to ensure high reflectivity and mechanical stability. The resulting air-to-water bridging reflector unit is suspended beneath the primary objective O1 via a custom-fabricated curved aluminum cantilever. With this configuration, we achieved a light sheet 5 mm in width, featuring a full-width-at-half-maximum (FWHM) beam waist of ~9.0 μm and a NA of ~0.035, corresponding to a confocal parameter (i.e., twice the Rayleigh range) of ~344 μm in water.

The excited planar fluorescence image, tilted by 60° in water relative to the optical axis of O1, is first refracted by the water-to-air interface and forms a virtual paraxial image tilted by 66.6° relative to O1’s optical axis (or 23.4° relative to O1’s focal plane). In other words, the epi-fluorescence image perceived by the dry primary objective O1 appears more oblique— albeit at the expense of increased spherical aberration. The primary detection path from O1 to O2 is configured for unity magnification, therefore the stationary intermediate image reconstructed in O2’s focal region is equally tilted by 23.4° relative to O2’s focal plane, thereby enhancing the fluorescence coupling efficiency into O3. In our implementation, we actually adopted a standard 4× objective with an NA of 0.14 as O3, given the high cost of large-format dichroic mirrors required for potential dual channel imaging.

Another side effect of the super-oblique light sheet generation approach is the non-lateral displacement of the natural sheet waist, as the inclined side-wall reflector mirrors the sheet waist onto an off-focal plane moving trace (see Supplementary Fig. S1). To compensate for this, we integrated an electrically tunable lens (ETL) to dynamically shift the sheet waist along the axial direction in synchrony with the sweeping motion of the illumination plane, so that the sheet waist stays nearly coplanar with the focal plane throughout the scan range. To minimize the impact of the ETL-induced diopter variation on the collimation and uniformity of the light sheet, the ETL was positioned conjugate to the back focal plane of the primary objective.

## 3. Experimental Results

### 3.1 Resolution characterization

To evaluate the resolution of the system, we imaged green fluorescent beads with a diameter of 500 nm embedded in a 1% agarose gel (Fig. 4A). The skew-corrected 3D volume data, measuring ~5.0 mm × ~2.9 mm in lateral dimension and ~0.30 mm in depth, was divided into 20 depth layers, each ~15 μm in thickness; then we randomly sampled 15 beads from each layer to calculate the per-layer FWHM statistics (Fig. 4B). The final average resolution was ~9.4 μm (x) × ~5.9 μm (y) × ~6.0 μm (z). The resolution is limited by the sheet thickness, spherical aberration associated with water-to-air interface, as well as other aberrations accumulated in the O1-SL1-TL1-SL2-TL2-O2 optical train.

**Fig. 4.**
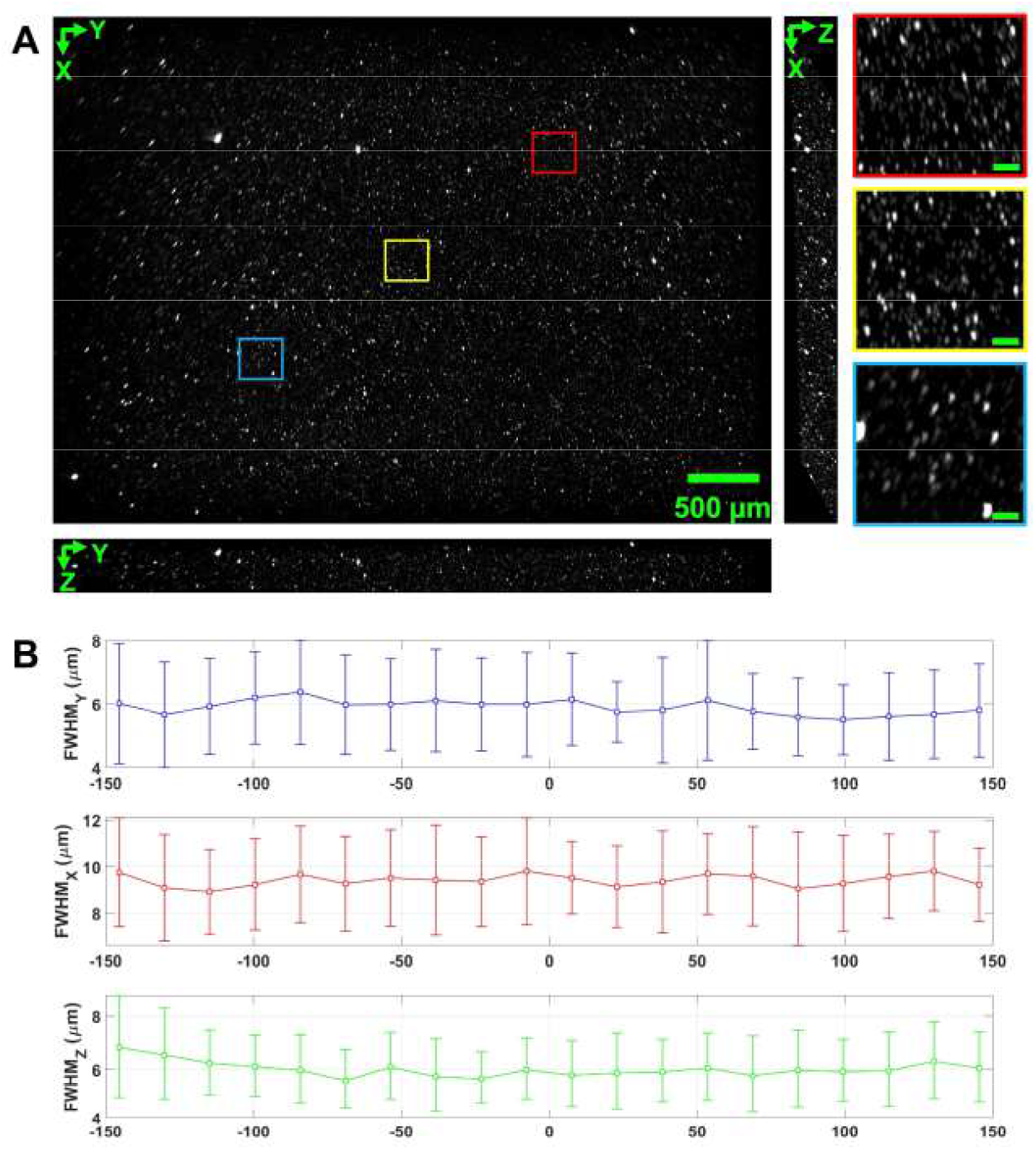
Resolution characterization. (A) Imaging results of 500-nm-diameter fluorescent beads embedded in 1% agarose gel, presented as maximal intensity projections (MIP) along the x-, y- and z-axis from the skew-corrected raw data spanning an *xyz*-FOV of 5.0 × 2.9 × 0.36 mm^3^. Insets on the right show close-up views of 3 boxed regions from the *xy*-plane MIP. Scale bar: 50 μm. (B) Spatial resolution statistics. Mean FWHM and standard deviation (calculated over 15 randomly selected beads per 15-μm-thick depth layer) are plotted against the central depth (in μm) of each layer relative to the natural focal plane of O1.

### 3.2 Whole body imaging of zebrafish larvae

In our experiments, we utilized a zebrafish larvae model with vascular endothelial cells labeled with GFP. The first experiment involved static imaging of the vascular endothelial cells across the entire body of a zebrafish. To verify that our system could achieve the desired field of view within biological tissue, we used zebrafish that had been reared for 8 days post-hatching and preserved in formalin, with a body length of approximately 4 mm. During imaging, the sample was embedded in agar, its body posture was not perfectly adjusted. To ensure full coverage of the body, the imaging depth reached nearly 1 mm, as seen in the side view (Fig. 5A). The imaging quality across different depths demonstrated the stability of our system in depth-resolved imaging. However, due to the reduced transparency of the head region, the imaging quality gradually degraded with increasing depth.

**Fig. 5.**
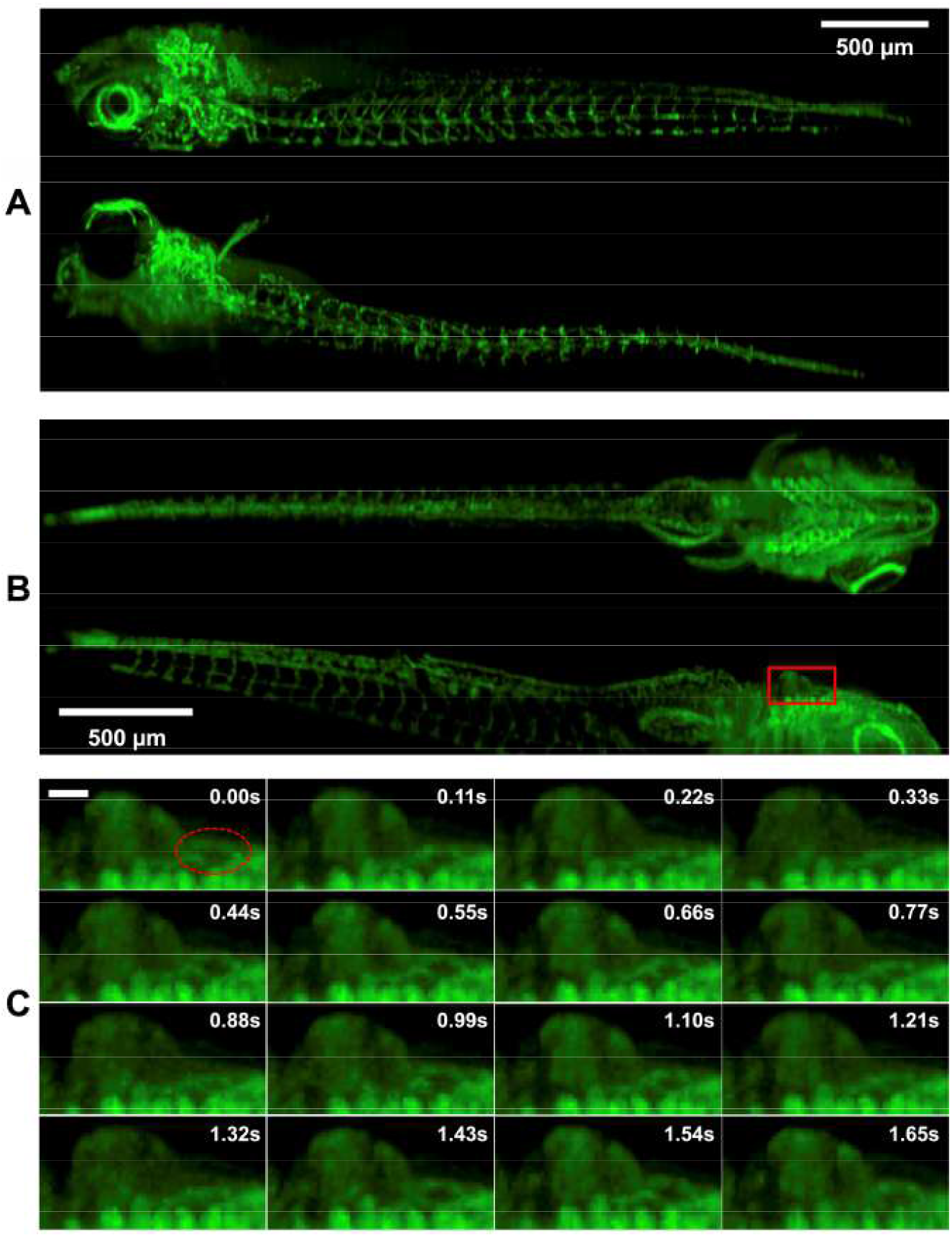
Whole body imaging of larval zebrafish. (A) Maximum intensity projections (top and side views) of a formaldehyde-fixed zebrafish larva embedded in agarose, with its vascular endothelial cells labeled by FLK1-GFP. (B) Imaging of a live zebrafish larva embedded in agarose. To capture cardiac dynamics, a shorter focal length tube lens was employed in the tertiary objective module, trading spatial resolution for a higher volumetric imaging rate of 9 Hz. (C) Close-up views of the cardiac region in the maximum intensity projection, displayed sequentially from left to right and top to bottom, with a time interval of 0.11 seconds. Scale bar: 50 μm.

The second experiment focused on dynamic imaging of live zebrafish. Since the zebrafish were immobilized in agar, their body movements were random, so we primarily observed their cardiac activity (Fig. 5B). We performed whole-body imaging at a low magnification and a volumetric rate of 9 Hz, successfully capturing the heartbeat (Fig. 5C). These experiments demonstrate the effectiveness of our rescanning strategy in mesoscopic imaging.

## 4. Discussion

In this work, we present a meso-SCAPE microscope designed to balance imaging performance, system complexity and cost-effectiveness. Rather than investing significant effort or relying on optical design professionals to customize a high-NA, mesoscopic FOV objective—and fabricating three such objectives as required by SCAPE—we built our system around commercially available large-FOV objectives with limited numerical aperture (NA). To enhance performance, we paired the low-NA primary objective (O1) with an easy-to-fabricate, air-to-water bridging reflector that enables the illumination plane to reach a steep oblique angle (~60° in water relative to the axis of O1), significantly surpassing the nominal NA of O1 (~0.5, or an aperture of 30° in air), while introducing minimal aberrations or engineering complexity. This RI-matched polymer-based trans-medium reflector shares its roots with the trapezoidal glass prism used in the light-sheet mirroring-based mesoscopic OPM design [29], yet our implementation is more compact, compatible with both upright and inverted imaging geometries, and supports flexible selection of polymer materials to match the RI of the various imaging targets.

Since the epi-fluorescence light goes through the polymer prism without reflecting off its sidewall, the optical reciprocity between the illumination and detection paths is disrupted, causing the natural descanning mechanism of the SCAPE microscope—fundamentally akin to that of a confocal microscope—to fail. To overcome this limitation, we developed a single-galvo-based, self-conjugating rescanning strategy that enables steering the illumination beam with an arbitrarily tunable optical deflection angle—no longer fixed to twice the mechanical rotation angle of the galvo mirror—to accommodate the modulated displacement of the super-oblique illumination plane, and eventually to restore descanned epi-fluorescence detection. The key principle involves introducing an imaging path that conjugates the galvo mirror to itself, such that the illumination beam, after its initially reflection, is routed back to the same galvo for a second scan (i.e., rescan). This self-conjugating rescanning strategy bears conceptual resemblance to the scan multiplier unit proposed for ultrafast laser scanning [31]. Notably, our strategy can be configured with an optical-to-mechanical angle ratio (OMAR) significantly greater than 1.0 (by adjusting the angular magnification of the self-conjugating imaging path), thereby achieving a given optical deflection with a much smaller mechanical scan angle and subsequently a higher galvo scanning frequency, and improving the throughput-speed product of a general scanning microscope [32].

The primary objective we chose (MVPLAPO 2XC, Olympus) supports a nominal FOV up to 17 mm in diameter for wide-field imaging [33], and a usable FOV of ~7-9 mm (with its rear aperture underfilled) [30]. However, in our prototype meso-SCAPE microscope, we observed that the optical aberrations grow prominent when the oblique illumination plane scans beyond 3.0 mm laterally. Through ZEMAX simulations and experimental characterization, we determined that the primary culprit lies in off-axis aberrations in both the illumination and collection optical paths. These aberrations manifest in two ways: 1) curling of the illumination light sheet, resulting in a non-planar intermediate image, and 2) field curvature, which worsens defocus-induced blur in the acquired intermediate image. To accommodate the 45-mm-diameter pupil of O1 (given by 45 mm EFL times 0.5 NA), both TL1 and SL1 employ Plössl lenses constructed from 3-inch achromats. The illumination beam and epi-fluorescence light pass through different off-axis regions of the SL1-TL1 relay system, making it challenging to simultaneously optimize both the illumination and detection paths—particularly at peripheral scan positions. In our current implementation, we have adopted a compromise design to balance resolution and lateral scan range. Further investigations are ongoing to search for improved Plössl lens combinations for SL1, TL1, SL2 and TL2, either relying entirely on commercial lenses or mixing off-the-shelf achromats with 1-2 custom lenses featuring optimized surface profiles [34, 35].

## 5. Methods

### Initial light sheet generation module

Illumination light from a 488 nm laser (OBIS 488LX, Coherent) is first expanded into an initial light sheet using a Powell lens (30° fan angle; LGL130, Thorlabs) in combination with an achromatic lens (RL_1_: EFL = 30 mm, GUO-145050, GU Optics). The resulting beam is then relayed onto the galvanometer mirror via a series of relay lens (RL_2_: EFL = 100 mm, GUO-145058, GU Optics; RL_3_ and RL_4_: EFL = 150 mm, GCL-010605, Daheng Optics). An ETL (EL-10-30-TC, Optotune) is positioned nearly the shared focal plane of RL_2_ and RL_3_, such that its varying diopter primarily modulates the axial position of the light sheet waist while minimally affecting the collimation and uniformity along sheet width direction. The ETL’s diopter is designed to vary from 8.3 m^-1^ to 20 m^-1^. To enable bi-directional shift of the sheet waist, we introduced a concave cylindrical lens (EFL = −75 mm, GCL-110313, Daheng Optics), pressed against the ETL housing, to offset the overall effective diopter range along the sheet width direction to approximately [−5.0 m^-1^, +6.7 m^-1^], corresponding to an EFL range of (−∞, −200 mm) ∪ (+150 mm, +∞). Such configuration enables axially shifting of the eventual super-oblique light sheet’s waist while keeping its waist nearly coplanar with O1’s focal plane across the desired scan range.

### Self-conjugating rescanning module

The pivotal point of the galvanometer scanner is imaged onto itself through two concatenated 4f-systems comprising four 2-in-diameter achromats (GL_1_: EFL = 150 mm, GCL-010616A, Daheng Optics; GL_2_, GL_3_ and GL_4_: GUO-145086, EFL = 100 mm, GU Optics). The entire self-conjugating optical imaging path is folded into a closed loop by three silver mirrors and a 2-in-diameter dichroic mirror (DM20-505LP, LBTEK) that reflects the excitation light towards the galvo scanner and transmits the epi-fluorescence (descanned by the galvo) to the secondary objective module. The incident light sheet is first reflected and scanned by the galvo scanner, and then relayed by the self-conjugating optical path back to the (still rotating) galvo scanner, where it gets reflected and scanned again (i.e., rescanned). Given the angular magnification of γ = 1.5 of the self-conjugating optical imaging path, the optical scan angle of the rescanned light sheet exiting the galvo is given by *θ* _ill_|_rescan_ = 2(1 − γ)*θ*_galvo_ = −*θ*_galvo_, yielding an OMAR of −1.

When the galvanometer is parked in its neutral position (i.e., at a 45° angle relative to the optical axes of both the O1 and O2 modules), the rescanned beam exits the galvo with an inclination of approximate 7.6° relative to the optical axis of the O1 module, leading to the illumination beam later exiting O1 parallel to O1’s optical axis with a lateral offset of 3.0 mm.

### Primary objective (O1) module

To match the optical throughput of the selected O1 (EFL = 45 mm, NA = 0.5), we built both scan lens SL_1_ and tube lens TL_1_ as Plössl lenses, each comprising two 3-in-diameter achromatic doublets, to minimize beam clipping and aberrations. The EFL of SL_1_ (88596 and 33925, Edmund Optics) and TL1 (88597 and 88598, Edmund Optics) are approximately 171.4 mm and 85.7 mm, respectively, yielding an angular magnification of −0.5 from the galvanometer to the back pupil of O1. As a result, the light sheet exiting O1—prior to interacting with the polymer prism—undergoes a lateral translation described by

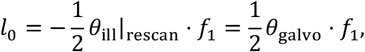

where counter-clockwise rotation and rightward translation are defined as positive.

The custom-built trapezoidal polymer prism, approximately 5.0 mm in thickness, features a 30-degree angled sidewall cured directly onto the silver-coated hypotenuse surface of a prism-shaped mirror. The top (air-facing) and bottom (sample-facing) surfaces are sealed with a ~170-μm-thick coverslip for protection. The light sheet exiting O1 enters the polymer prism perpendicularly, reflects off the angled sidewall, and exits the bottom surface at an oblique angle of about 60° relative to the optical axis of O1. Compared to the pre-reflection light sheet, this reflected super-oblique light sheet laterally translates at double the speed but in the opposite direction. This results in a net lateral displacement of

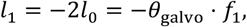

which satisfies the condition for descanned epi-fluorescence detection (as detailed shortly).

The entire air-to-water bridging reflector unit (i.e., the trapezoidal polymer prism and the silver mirror) is mounted within a custom 3D-printed holder and suspended beneath O1 using a precision-machined curved aluminum cantilever. Through careful mechanical alignment, we positioned the bottom coverslip ~1250 μm below O1’s natural focal plane since the epi-fluorescence image formed by the water-to-air interface appears closer (i.e., moving upwards). Furthermore, proper adjustment of the ETL’s diopter enabled the light sheet waist to be brought into exact alignment with the focal plane.

### Secondary objective (O2) module

Epi-fluorescence emitted from the illumination plane passes directly through the polymer trapezoid, without reflecting off its angled inner sidewall, and propagates all the way back through O1, TL_1_, and SL_1_ towards the rotated galvanometer mirror. Given the lateral displacement of the super-oblique light sheet (*l*_1_ = −*θ*_galvo_ · *f*_1_ as derived previously), the returning epi-fluorescence rays—prior to reflection by the galvanometer—are deflected by an angle

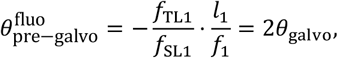

with counter-clockwise rotation (or deflection) defined as positive. Since the galvanometer mirror itself has rotated by *θ*_galvo_, the post-galvo reflection epi-fluorescence rays are effectively descanned, resulting in the formation of a stationary intermediate image at the focal region of O2 (see Supplementary Note S1 for detailed analysis).

We selected O2 to be identical to O1, and paired it with a 60-mm-diameter achromatic scan lens (SL_2_; EFL = 100 mm, GUO-145097, GU Optics) and a 3-inch-diameter achromatic tube lens (TL_2_; EFL = 200 mm, 88596, Edmund Optics). This setup ensures matched optical throughput while providing ample space for deploying the self-conjugating rescanning module (as shown in Fig. 2C). This configuration establishes a unity lateral magnification from the focal space of O1 to that of O2—or more precisely, between the virtual image formed by the polymer-to-air interface beneath O1 and the stationary intermediate image formed in the focal region of O2. Factoring in the 1.0× lateral and 0.75× axial magnification of the polymer-to-air interface, the resulting intermediate image is tilted by approximately 23.4° relative to the focal plane of O2, thereby enhancing the fluorescence coupling efficiency into O3.

### Tertiary objective (O3) module

Given the size of the intermediate image, we can in principle select O3 to be identical to O1 and O2, yielding a fluorescence coupling efficiency of about 52.3%; however, this results in a large pupil size (approximately 45 mm in dimension), rendering it challenging and costly to design and fabricate a matching dual-channel image splitter. Therefore, for proof-of-concept purpose, a cost-effective 5× commercial objective (EFL = 40 mm) was chosen as O3 with an ample FOV but a much lower collection NA of 0.14. Thanks to the super-oblique illumination light sheet and intermediate image, such low-NA objective still affords considerable fluorescence efficiency. O3 was paired with a tube lens (TL_3_) of appropriate EFLs (e.g., 135 mm EFL for resolution characterization, or 50 mm EFL for rapid volumetric imaging.

To accommodate the expanded FOV and increased imaging throughput, we selected an sCMOS camera with features 3200×3200 pixels and a peak quantum efficiency of 95% (Kinetix, Teledyne Photometrics). For a typical imaging depth of ~500 μm—corresponding to ~1000 μm along the z’ direction—we allocate at least 400 rows on the sensor to satisfy the Nyquist sampling criterion. In practical applications, fewer rows can be used (i.e., sub-Nyquist sampling) to achieve higher frame rate and faster volumetric imaging.

### Synchronization and descan calibration

The galvo scanner, the ETL, and the sCMOS camera are orchestrated via a DAQ card (PCIe-6321, National Instruments), which is in turn configured and controlled by a home-developed GUI software written in C/C++ and Qt. At the start of each volume scanning task, the user needs to adjust the sample in preview mode and confirms the FOV to use. Then the control software will configure the DAQ card to output appropriate voltage signals for the galvanometer, the ETL, and the sCMOS camera, respectively according to the user-set volume scanning parameters. These three signals are synchronously (re-)triggered by a TTL clock output from a counter on the DAQ card for reproducible initiation of volumetric data acquisition.

Successful descanning requires perfect matching between the optical scan angle of the self-conjugating rescanning module and the modulated lateral displacement of the super-oblique illumination plane. For this purpose, we need to carefully adjust the orientation of the trans-medium bridge reflector prior to imaging experiments. The current calibration procedure involves scanning an in-house fabricated thin fluorescent reference film (sealed and protected with a cover glass), while monitoring the line image acquired by the properly aligned tertiary module. Calibration completion is confirmed when the line image remains stationary throughout galvo scanning.

### Data pre-processing

We use MATLAB to preprocess the collected data. First, we determine the pixel spacing in each dimension of the image stack. ΔX corresponds to the sampling spacing in the scanning direction, ΔY represents the actual spatial sampling spacing of the camera pixels on the sample, and the ΔZ represents the inter-layer distance in the depth direction. Then, we perform a shear transformation on the volume data in the X direction to reconstruct it into volume data with the original scale.

## 6. Supplementary Materials

### 6.1 Supplementary Note S1

To analyze the deflection angle of both incident and emission light (or rays), we take as neutral reference the configuration A in Fig. 2A, where the reflected super-oblique illumination plane crosses the front focal point of O1. Then, angles of galvanometer rotation and (illumination or epi-fluorescence) beam deflection referred to in the following analysis are all relative to their counterparts in this neutral reference, with counter-clockwise defined as the positive direction.

With the galvo mirror mechanically scanning/rotating by *θ*_galvo_, shown as configuration B in Fig. 2A, the optical scan angle of reflected illumination beam (or light sheet) exiting the galvo mirror equals *θ*_ill_ = 2*θ*_galvo_. Going through the 4f-system comprising a scan lens SL_1_ and a tube lens TL_1_, with their effective focal lengths denoted as *f*_S_ and *f*_T_ respectively, the illumination plane eventually exits O1 and reflects off the reflector surface of the trapezoidal prism, turning into a super-oblique illumination plane that displaces to position B in the focal plane. With rightward displacement defined as positive, the lateral translation of the super-oblique illumination plane could be derived as

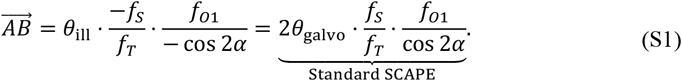

Epi-fluorescence light (light blue shadow) emanating from the illuminated plane, collected by O1 and propagating through the tube lens and the scan lens, impinges onto the galvo with a deflection angle (light blue shadow, relative the counterpart in configuration A, with still counter-clockwise defined as positive) of

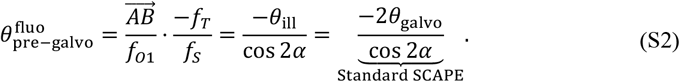

The deflection of the epi-fluorescence light reflected off the galvo in the current configuration B, relative to the reflected epi-fluorescence light in configuration A, results from the linear superposition of two effects. First, for a fixed galvo mirror, when the incident beam itself deflects by *θ* _incident_, the reflected beam will deflect along an inverse angular direction, i.e., by −*θ*_incident_; secondly, for a given incident beam, when the galvo scans by *θ*_galvo_, the reflected beam will scan along the same angular direction by 2 · *θ*_galvo_. Therefore, the total relative deflection of the reflected epi-fluorescence light amounts to

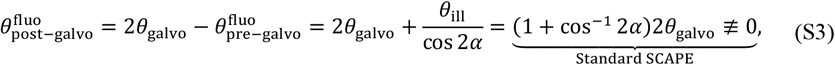

which implies that the descanning mechanism fails for a super-oblique illumination plane when using the galvanometer scanner in a standard way.

In light of this, we proposed a self-conjugating rescanning module which breaks the *θ*_ill_ = 2*θ*_galvo_ convention. Instead, the optical-to-mechanical scan angle ratio (OMAR), defined as *b* ≐ *θ* _ill_/*θ*_galvo_ can be flexibly customized. One can easily see that by setting *b* = −2 cos 2*α* and substituting it into Eq. (S3), the right-hand side evolves to

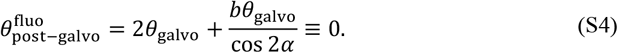

A conceptual schematic of the self-conjugating rescanning module we proposed to achieve such customizable OMAR is illustrated in Fig. 2B. The core is to introduce an imaging path with an angular magnification factor γ that maps the rotational center of the galvanometer mirror back onto itself—hence the term “self-conjugating.” In this configuration, the illumination beam is first reflected by the galvanometer mirror, then re-directed through the imaging path, and subsequently re-incident on the same mirror. As a result, the galvanometer’s rotation modulates the beam twice—constituting the so-called “rescanning.”

Following the sign convention used in previous analysis, after the first reflection, the illumination beam is deflected by an angle of *θ*_first_ = 2*θ*_galvo_. When the beam re-enters the galvanometer mirror via the self-conjugating imaging path with angular magnification γ, its deflection angle becomes *θ*_self-conj_= γ · *θ*_first_ = 2γ · *θ*_galvo_.

After the second reflection off the rotated galvanometer, the total deflection angle of the exiting beam—following the same logic when deriving Eq. (S3)—is given by

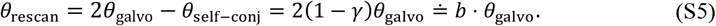

By appropriately designing the angular magnification γ of the self-conjugating imaging path, one can flexibly customize the desired OMAR *b* = 2(1 − γ), thereby meeting the requirements for descanned detection.

### 6.2 Supplementary Note S2

Another issue associated with the super-oblique light sheet generation strategy is that the natural waist of the super-oblique light sheet, when scanned, no longer stays coplanar with O1’s focal plane. Instead, as shown in Figure S1A, due to the mirroring effect of the inclined reflecting surface, the waist of the reflected light sheet moves within the mirror plane of the focal plane. This results in the thinnest (and also optimal) portion of the reflected illumination plane deviating from the focal plane that affords in general the optimal detection PSF for the detection path, leading to significant degradation of the final image quality.

**Figure S1.**
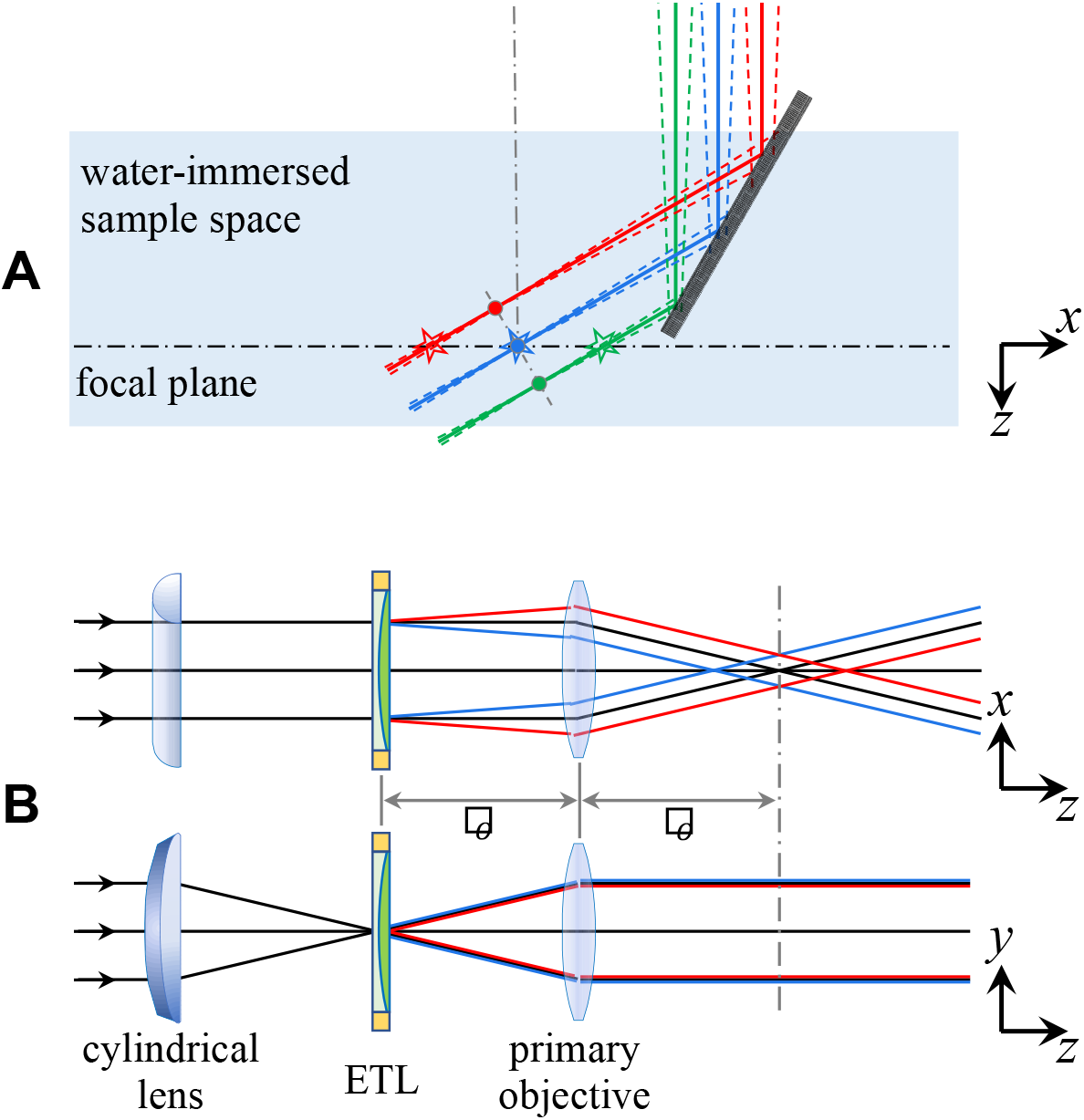
Mechanism and design of ETL-based axial sheet waist position compensation. (A) Scanning trajectories of the oblique light sheet waist before (circles) and after (pentagrams) correction, with red, blue, and green colors representing different scanning positions. (B) Working principle and fundamental optical path for minimizing light sheet deformation through axial waist position adjustment.

To address this, we implemented dynamic compensation of the axial displacement of the light sheet waist based on an electrically tunable lens (ETL). The fundamental principle is to adopt a series of cascaded 4f-system to conjugate the ETL to the rear focal plane of the primary objective, thereby exerting extra convergence or divergence to the wavefront and subsequently altering the axial position of the light sheet waist.

To minimize the impact of ETL focal power adjustment on the overall shape and illumination uniformity of the modulated light sheet, the cylindrical lens employed to launch the initial light sheet should be positioned upstream of the ETL, and the waist of initial light sheet, the ETL, and O1’s pupil plane should be conjugated to each other (the three planes are drawn as coincident for simplicity in Fig. S1B). In this way, while adjusting the ETL’s focal power, the overall optical power of the ETL-O1 composite lens restricted in the *xz*-plane remains unchanged, with only the principal plane shifting axially. This allows the post-O1 light sheet waist to move forward or backward and leaves the light sheet’s numerical aperture (NA) unchanged. Moreover, in the yz plane, as incident rays bent by the cylindrical lens converge approximately at the principal plane of the ETL, the direction of exiting rays stays roughly independent of the ETL’s focal power. Consequently, the final light sheet remains collimated in its width direction.

### 6.3 Supplementary Note S3

**Figure S2.**
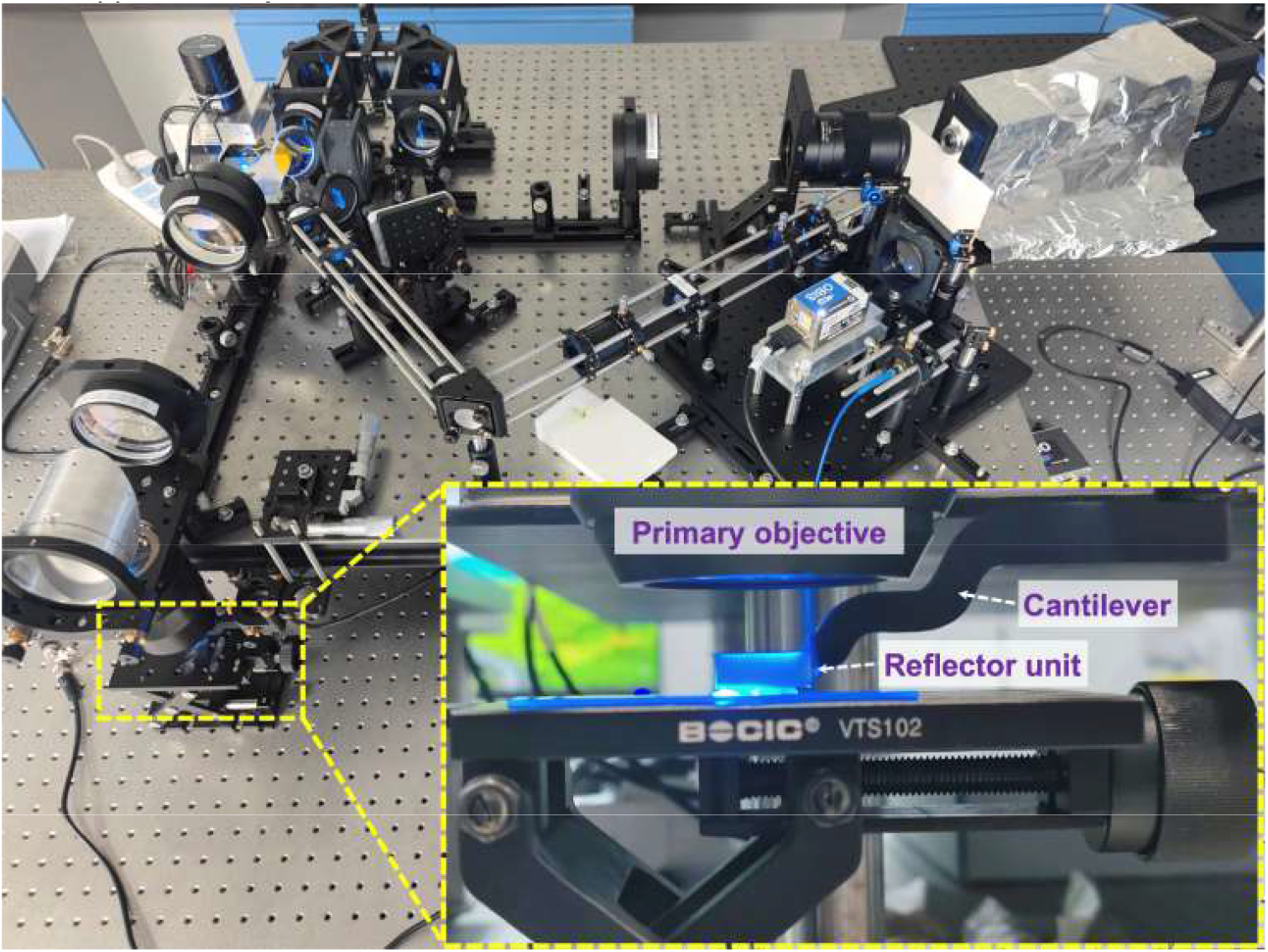
Photograph of the prototype meso-SCAPE microscope.

## Notes

### Competing Interest Statement

The authors have declared no competing interest.

